# The role of vimentin-nuclear interactions in persistent cell motility through confined spaces

**DOI:** 10.1101/2021.03.16.435670

**Authors:** Sarthak Gupta, Alison E. Patteson, J. M. Schwarz

## Abstract

The ability of cells to move through small spaces depends on the mechanical properties of the cellular cytoskeleton and on nuclear deformability. In mammalian cells, the cytoskeleton is comprised of three interacting, semi-flexible polymer networks: actin, microtubules, and intermediate filaments (IF). Recent experiments of mouse embryonic fibroblasts with and without vimentin have shown that the IF vimentin plays a role in confined cell motility. We, therefore, develop a minimal model of cells moving through confined geometries that effectively includes all three types of cytoskeletal filaments with a cell consisting of an actomyosin cortex and a deformable cell nucleus and mechanical connections between the two cortices—the outer actomyosin one and the inner nuclear one. By decreasing the amount of vimentin, we find that the cell speed is typically faster for vimentin-null cells as compared to cells with vimentin. Vimentin-null cells also contain more deformed nuclei in confinement. Finally, vimentin affects nucleus positioning within the cell. By positing that as the nucleus position deviates further from the center of mass of the cell, microtubules become more oriented in a particular direction to enhance cell persistence or polarity, we show that vimentin-nulls are more persistent than vimentin-full cells. The enhanced persistence indicates that the vimentin-null cells are more subjugated by the confinement since their internal polarization mechanism that depends on cross-talk of the centrosome with the nucleus and other cytoskeletal connections is diminished. In other words, the vimentin-null cells rely more heavily on external cues. Our modeling results present a quantitative interpretation for recent experiments and have implications for understanding the role of vimentin in the epithelial-mesenchymal transition.

## I. INTRODUCTION

Cell migration is a fundamental process that contributes to building and maintaining tissue. To be able to migrate, the cell cytoskeleton, which is comprised of three dynamic polymeric systems: F-actin, microtubules, and intermediate filaments (IFs), generates forces. While actin and microtubules are more studied cytoskeletal filaments, intermediate filaments (IFs) also play a role in a range of cell and tissue functions [1–3]. We will focus here on the IF protein known as vimentin. Vimentin is an IF protein whose expression correlates with *in vivo* cell motility [4, 5] behaviors involved in wound healing [6, 7] and cancer metastasis [8, 9], and, yet, its role in three-dimensional cell migration is poorly understood.

Typical modeling of cells migrating on surfaces involves the actin cortex with actin polymerization driving the leading edge of the cell, adhesion created at the leading edge, adhesion disassembled at the rear, and actin-myosin contraction at the rear as well so that the rear keeps up with the leading edge [10]. More recent surface models exist and do not involve the nucleus [11–14]. And while such models made progress accurately represent cell speed, predicting cell direction has been more elusive [15]. In an effort to understand the polarity of a migrating cell on a surface, some models include both actin and microtubules [16, 17]. It has also been reported that the nucleus has a crucial role in cell polarity in three dimensions but is dispensable for one and two-dimensional motion [18]. Two-dimensional experiments indicate that the centrosome, the main microtubule-organizing center, coordinates the direction of cell migration and is typically positioned between the nucleus and the leading edge of the cell [19].

While surface cell motility experimental and theoretical studies are plentiful, confined cell motility may indeed more representative of *in vivo* cell motility where cells move in the presence of other cells and/or in the presence of extracellular matrix (see. Figs 1(a) and 1(b)) [20]. Emerging experimental studies show that the structure and dynamics of the cytoskeleton in three-dimensional motility differ from those for cells on surfaces [21]. In confined settings, actin tends to accumulate at the cell cortex [22] and microtubules align with the direction of the confining track [23]. However, recent studies have shown that in three-dimensional settings the centrosome displays an increased probability to be near the rear of the cell [24]. When cells change direction in a narrow track, the centrosome repositions, moving from one side of the cell to the other, to repolarize the cell by developing a new trailing edge of the cell and setting up the polar direction of the cell. Interestingly, the nucleus appears to be decoupled from this phenomenon in that the removal of the cell nucleus does not alter the centrosome repositioning in most cells [24]. Though for high enough confinement [25], the cell nucleus will presumably play a more dominant role in the repositioning of the centrosome as it deforms. Prior cell motility models in strong confinement, therefore, focus on the nucleus [26–30].

**FIG. 1.**
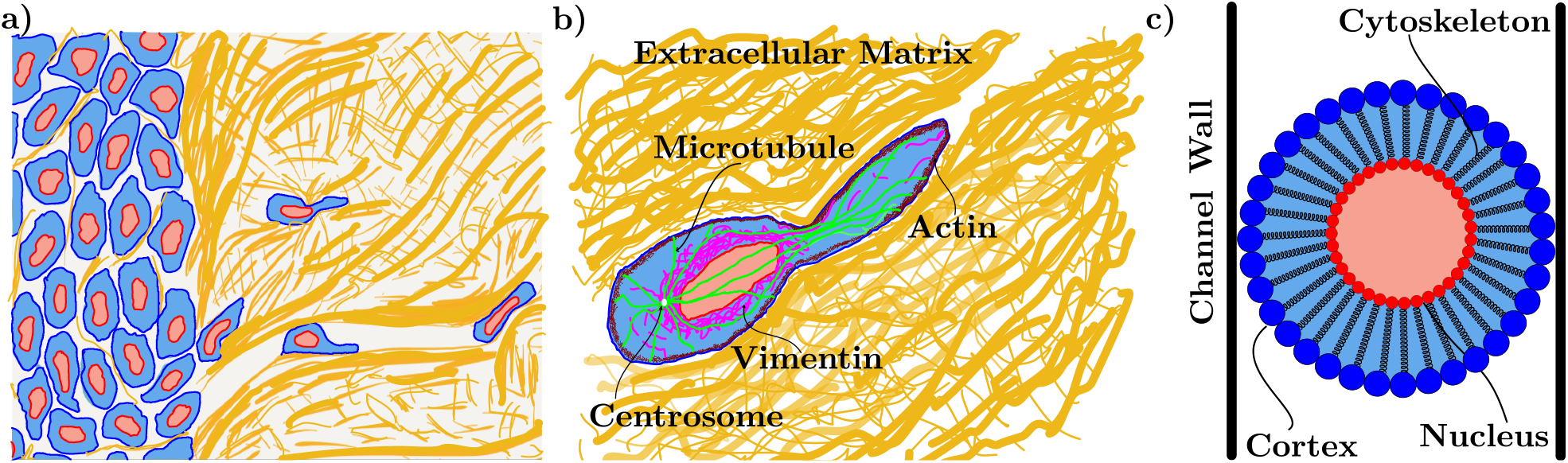
Modeling confined cell motility: (a) Cells move through confining environments due to the extracellular matrix (in gold) and other cells. (b) Schematic of an individual cell with a cytoskeletal network containing actin, microtubules, and intermediate filaments, a nucleus, and a centrosome connected to the nucleus via the protein dynein. (c) A simulation model for the cell: The cortex is made up of blue monomers connected with springs, the nucleus is also made up of red monomers with springs connecting them. The bulk cytoskeletal network is simplified and modeled as springs connecting the nucleus and the cortex. The cell and nucleus contain cytoplasmic and nucleoplasmic material, each of which is modeled as area springs.

Given the roles of actin, microtubules, and centro-some positioning in confined cell motility, one is naturally led to ask about the role of vimentin, which couples to both actin and microtubules and forms a cage around the cell nucleus [31]. Recent experiments by one of us [32], and others [33, 34], highlight new roles of vimentin in mediating cell speed and cell persistence in three-dimensional confining environments. Specifically, when comparing the motion of wild-type mouse embryonic fibroblasts (mEFs) with their vimentin-null counterparts moving through three-dimensional micro-fluidic channels, the loss of vimentin increases cell motility and increases rates of nuclear damage in the form of nuclear envelope rupture and DNA breaks. Moreover, unlike in unconfined motility, the loss of vimentin increases the spontaneous persistent migration of cells through 3D micro-channels and 3D collagen matrixes [35]. Additional experiments demonstrate that vimentin may also play a more active role in persistent cell motility via its interactions with actin stress fiber [36] formation and microtubule positioning [37].

Inspired by these experiments, we seek to quantitatively understand the role of vimentin in confined cell motility via a computational model for the first time. To do so, we develop a minimal cell motility model incorporating actin, microtubules, and vimentin. The starting point of the model is the hypothesis that vimentin plays a distinct role in mediating forces between the actomyosin cortex and the nucleus [38, 39]. Our model, therefore, contains both an actomyosin cortex and a nucleus, whose interaction via a set of linker springs is strengthened by the presence of vimentin. We also propose a new mechanism for cell polarity for motility in which vimentin plays an important role. As the cell moves through the model micro-channel, we will quantitatively show how vimentin modulates cell speed, nuclear shape, dynamics, and cell persistence. In addition to quantitative comparison with prior experiments, we pose new predictions for the role of vimentin in confined cell motility more generally. In particular, we address the upregulation of vimentin typically found in the epithelial-mesenchymal transition [5, 40].

## II. MODEL

### The players

We begin with the cellular cytoskeleton, composed of actin, microtubules, and intermediate filaments. The persistence length of actin filaments is smaller than microtubules but larger than intermediate filaments [41]. Myosin motors exert forces on actin filaments to reconfigure them. Many actin filaments and myosin motors reside in proximity to the cell membrane to form the actomyosin cortex. The actomyosin cortex is an important piece of the cell motility machinery as its reconfiguring drives cell motion. Microtubules typically originate from the microtubule-organizing center, or centrosome, and have a crucial role in cell polarity as they as required to generate traction forces [16]. Microtubules also have a role in controlling cell shape as they typically span the entire cell and are the stiffest cytoskeletal filaments [41]. Vimentin filaments exist as a cage or mesh structure around the nucleus and are also present in the cytoskeleton in fibrous form as can be seen in Fig. 1(b) [31, 38]. Studies show that vimentin provides structural integrity to the cell [25, 42] and also provides organization for the other cellular parts [43, 44].

The nucleus is typically the largest, and stiffest, organelle in the cell, yet it is still deformable. Recent studies showed a cell under a highly confined environment, and thereby, the nucleus is so squeezed such that DNA becomes damaged [25]. Cells cannot move through a particular confining geometry if the nucleus cannot do so [45]. Therefore, the nucleus is also an important player in confined cell motility.

These individual players do not act independently of each other. Major cytoskeletal network actin, microtubules, and intermediate filaments are also highly interconnected with each other[46]. For instance, vimentin and actin directly interact with each other via the tail domain of vimentin [47]. Moreover, plectin is a major crosslinker among all three types of filaments [48, 49]. The interconnected cytoskeletal networks and the nucleus are also interconnected among themselves via different proteins, motors, crosslinkers, etc. The inner nuclear membrane contains a protein network consists of Lamin A/C, which affects the size and shape of the nucleus [50] and disturbing them can result in defective nuclear mechanics [51]. LINC complexes made of nesprins and Sun proteins act as connecting bridges between nuclueoskeleton and actin network in cytoplasm [52]. Microtubules are also joined to the nucleus via kinesin-1 which talks to nesprin-4 [53]. Intermediate filaments are connected via plectin which connects to nesprin-3, which is joined with nucleus [54]. These LINC complexes also mediate intercellular forces from the cytoskeleton to the nucleus and affect the position and shape of the nucleus in the cell [55]. Disruption of these nucleus-cytoskeleton links also leads to impairment of 3D cell migration [56]. Therefore, the nuclear envelope, or cortex, consists of inner nuclear lamins and outer vimentin.

### Spring network

Given the complexity of such interactions, to model the interplay between cell mechanics, shape, and motility, we took a reductionist approach and simplified this highly tangled picture to a two-dimensional network of harmonic springs with an outer ring of springs representing the cellular cortex and an inner ring of springs representing the nuclear envelope, and harmonic springs connecting the cellular cortex with the nuclear cortex (see Fig. 1(c)). Our model is a two-dimensional cross-section of the three-dimensional system. As we can see from the schematic, each cell cortex monomer is joined to each nuclear cortex monomer via a linker spring. Each spring type, the cellular cortex, the nuclear cortex, and the interacting bulk linker spring, has its own stiffness with the respective potential energies 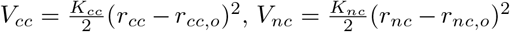, 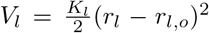 respectively, with *r_cc_* denoting the distance between the centers of neighboring cell cortex monomers, for example, and *r_cc,o_* represents the rest length of the spring. We also include potentials in the form of two area springs, 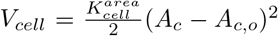 and 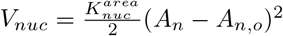, where *A_c_* and *A_n_* denote the areas of the cell and nucleus, respectively. The two area springs prevent the cell and the nucleus from collapsing as the cell becomes increasingly more confined.

How do we explore the role of vimentin in such a mechanical model, particularly when considering wild-type fibroblasts versus their vimentin-null counterparts? Since vimentin-null cells are softer than wild-type cells [42] and exhibit more DNA damage [25], to capture both cell lines we change the stiffness of the cytoplasmic/linker springs K_l_ and nucleus area springs K_nuc_. Since removing vimentin does not significantly affect the cortical stiffness [42], we do not alter the cortical spring constant among two cell lines.

The cell also interacts with its confining walls which are considered to be adhesive due to fibronectin or collagen I or some other kind of protein that is usually coated inside the channels. Specifically, the micro-channel is modeled as two lines of wall monomers fixed in place that interact with the cell cortex monomers via adhesion. In previous studies, adhesion to the surface is modeled as catch and slip bonds [57–59], where the cell forms a bond to the substrate, and then as it moves those connections peel off. We, therefore, model the adhesion interaction with the Weeks-Chandler-Anderson potential, or

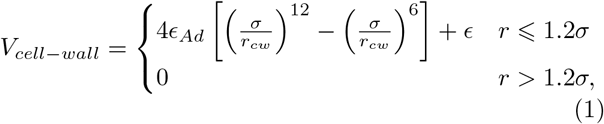

where *ϵ* quantifies the adhesion strength and the minimum of the potential is located at 2^1/6^*σ*. While the potential is purely repulsive, there is an attractive component (with a minimum) and a repulsive component. The minimum of the potential thus represents a typical integrin length [60] We did not vary the adhesion strength between the two cell lines.

### Dynamics

As the cell moves, actin is polymerized at the leading edge of the cell to translate the cell in a particular direction with microtubules setting the direction[61]. We model actin polymerization via an active force, **F**_*a*_. The active force is present for half cortex of the cell, the leading edge half, and has a magnitude *F_a_*. The direction of **F**_*a*_ is initially chosen to be towards the opening of the micro-channel, which determines the leading edge half—its polarization direction. There are small fluctuations in the direction of **F**_*a*_ as it moves through the micro-channel. In addition to adhering to the wall, it has been observed that in confined environment experiments, actin bundles start forming at the edges of the cell where it is interacting with the walls [62, 63]. We are proposing that due to this interaction with the wall, the cell generates forces to enhance motility in the direction in which it is polarized. Therefore, any cell cortex monomer with some proximity of a wall monomer exerts an additional force, **F**_*w*_ in the direction of the leading edge.

Now that we have detailed the forces involved, here is the equation of motion for each cell monomer of type *i* at position **r**_*i*_:

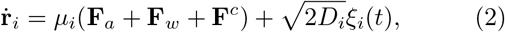

where **F**_*a*_ is an active force representing the actin forces at the front end of the cell in the direction of motion and **F**_*w*_ denotes the force generated at the wall in the direction of the leading edge. Finally, **F**^*c*^ represents the conservative forces in the system, or **F**^*c*^ = **F**_*cc*_ + **F**_*nc*_ + **F**_*l*_ + **F**_*cell*_ + **F**_*nuc*_ + **F**_*cell*−*wall*_), which are the two-body springs, the area springs, both modeling the mechanics of the cell, and the adhesion force between the cell and the confinement (wall). For the parameters varied in the simulations, see Table 1. We use simulation units defined as unit simulation length equal to *μ*m, unit simulation time is equal to *sec* and unit simulation force is *nN*. As for other parameters, D_cc_ = 0.02 *μ*m^2^*/s*, D_nc_ = 0.04 *μ*m^2^*/s* and *μ*_cc_ = 0.01 *μ*m nN/s, *μ*_nc_ = 0.02 *μ*m nN/s for both cell lines. The respective diameter of the monomers are *σ_cc_* = *σ_cw_* = 2 *μm*, *σ_nc_* = 1 *μm*, all neighbouring springs in actomyosin cortex and nuclear cortex is 2 *μm* and 1 *μm* respectively and linker spring length is 5.73 */mum*. Our simulations used 36 monomers in the both cortices, 74 monomers to simulate each side of the straight part of the wall, and 7 monomers for each side of the slanted channel entry and exit. To iterate Eq. 2, we use the Euler-Marayuma method.

**Table I.**
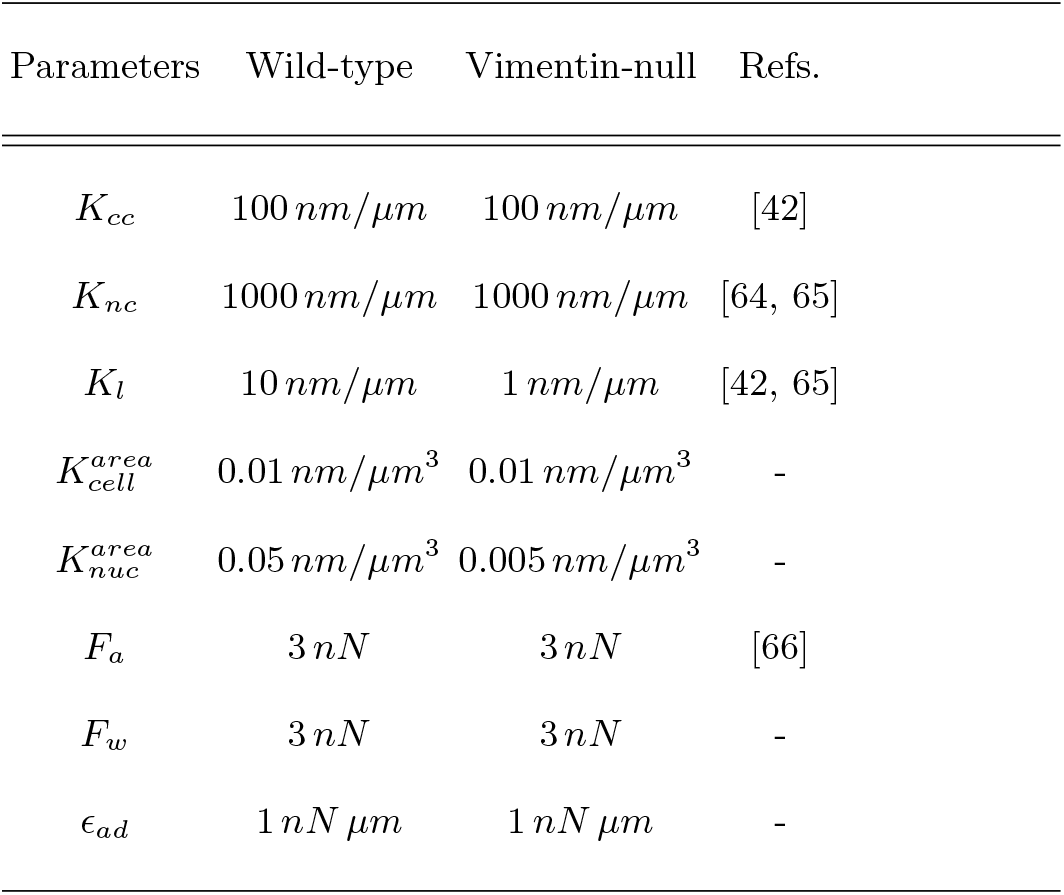
Table of parameters used, unless otherwise specified, and kPa is converted nm*/μ*m for the cortex, nucleus, and linker stiffnesses using a relevant length scale.

### Polarity-mechanism

If one allows for large fluctuations in the direction of **F**_*a*_, the probability is enhanced of the cell turning around in the channel. Recent experiments in two-dimensional motility suggest that removing the centrosome induces microtubules to grow symmetrically in all directions and, as a result, the cell forms lamellae in many directions and, therefore, loses its polarization [67]. In a 3D setting, the centrosome is typically found to be posterior of the cell, and microtubules are oriented in the direction of motion (for certain cell types) [68]. Therefore, the position of the centrosome sets the polarity of the cell by defining the tail/back of the cell during migration since the change in cell direction results only after the centrosome moves to the new posterior side of the cell [69]. Cell length is also found to be correlated with the time required for cells to change the direction, which is the time the centrosome takes to move to the other side of the cell to define a new tail [24]. Therefore, we posit that the more the centrosome is located away from the cell center of a crawling cell, the more biased cell migration becomes. Thus, the centrosome is essential for the preservation of polarized cell morphology.

While there is no explicit centrosome in our model, the centrosome is also connected to the nuclear outer membrane by a protein emerin [70]. Given the strong coupling between the nucleus and the centrosome, we effectively include a centrosome and a direction of cell polarity as determined by the direction of microtubule polymerization. To do so, we define **d** representing the difference between the center of mass of the cell and the center of mass of the nucleus. Its angle is measured from the positive *x*-axis. For incorporating the centrosome role in motility, we define the following empirical equation:

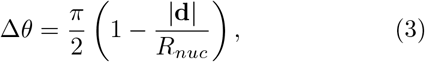

where *R_nuc_* denotes the radius of the nucleus. With this empirical equation, we define the bounds on the direction a cell can take. We take the nuclear cortex-cellular cortex (NC-CC) axis as our reference axis and for the upper bound and lower bound we can add and subtract Δ*θ* from this axis. See Fig. 4(a). Then, we choose a random angle under these bounds and that represents the direction of the nuclear, or centrosome, axis given the strong coupling between the two, which then drives the direction of migration.

What is the timescale for changing Δ*θ*? We assume that vimentin acts as a template for microtubules in the cell on a time scale of 10-20 minutes since (1) vimentin has a slow turnover rate than microtubules and (2) microtubules in vimentin-null cells show less orientation than in wild-type cells [37]. Therefore, this polarity mechanism repeats itself in every 15 min. as on average this is the time after which vimentin restructures. We have used the initial radius of the nucleus as a normalization constant in Eq. 3. Note that there is no memory of the previously taken direction. The NC-CC axis is, again, taken as a reference after each 15 min. and depending on the distance of the nucleus from the cell cortex, Δ*θ* is calculated, and eventually, a new angle is again chosen randomly from within the bounds. Note that the cell does not exactly follow the NC-CC axis for cell migration due to the intracellular dynamics and fluctuations in the cell.

### Analysis

We measure the average speed of the center of mass of the cell while it is in the channel. This cell speed average is then averaged over approximately 1000 realizations for each channel width. We also measure the persistence in the motility as defined as the ratio of the path length of the center of mass of the cell divided by the length of the channel. Flux is defined as the fraction of cells that exit the channel on the side different from the entry side. The circularity of the nucleus *C* is defined as 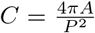, where *A* is the area of the cell and *P* is its perimeter. When *C* = 1, the nucleus is a circle. The time-averaged energy of the cell is calculated by taking the ensemble average of energy contributions from the conservative potentials. The initial cell energy for the conservative potentials is zero. The diffusion constant is extracted from the mean-squared-displacement of the cells as a function of time.

## III. RESULTS

### A. Vimentin affects optimal cell speed and its location as a function of channel width

Unlike 2D, wild-type cells move more slowly than vimentin-null cells in 3D channels [32]. See Fig. 2. Is this reduction in motility due to a decrease in cell speed and/or a decrease in persistence? Let us first address cell speed. For the wild-type cells, we observe that as the channel width decreases, the cell speed is non-monotonic. How does such a trend emerge? As the channels become narrower, the cell’s cortex increases its contact with the wall. This increase in contact generates more driving force to increase cell speed. However, this trend is competing with the deformability of the cell. As the channel width becomes even narrower, the linker springs become even more deformed (more compressed) and so these springs act to increase the effective adhesion to the wall given that unbinding to the wall is driven by a distance threshold. This increased adhesion time leads to a slower cell speed. Given the two competing trends—-contact with the wall increasing the driving force, yet also increasing the adhesion time as the cells become increasingly more deformed, one expects a nonmonotonic trend in the cell speed as a function of the channel width. We can modulate this competition by either increasing the driving force or the adhesion strength. If one increases the driving force, either by increasing the active force or the wall force, not only will the optimal speed increase, the peak will broaden towards smaller channel widths. See Supplemental Fig. S1. If one decreases the adhesion strength, a similar effect occurs. See Supplemental Fig. S2.

**FIG. 2.**
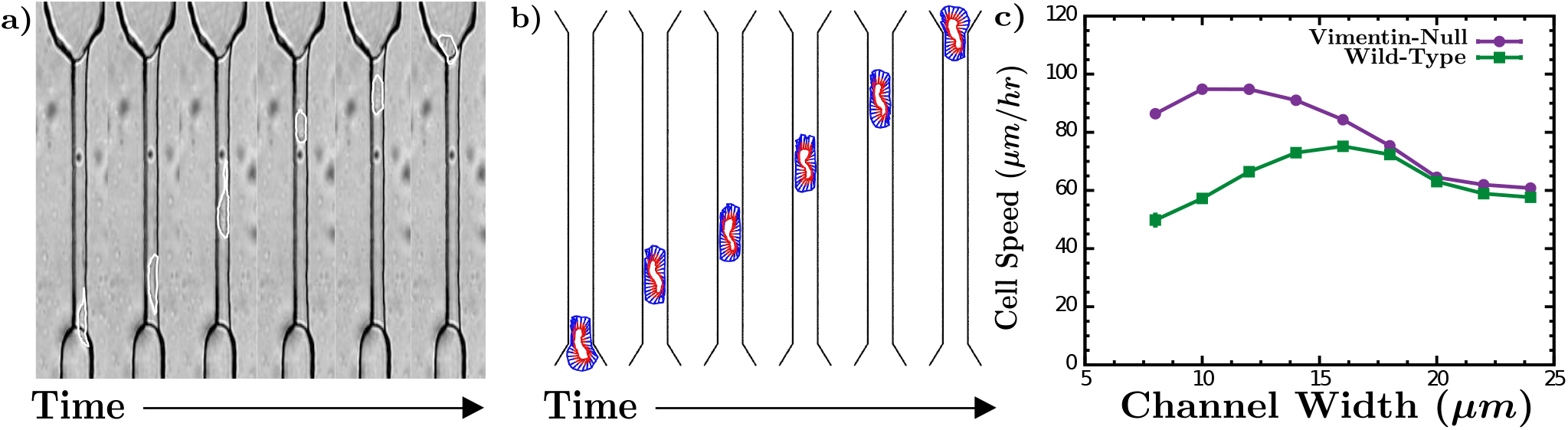
Model quantitatively agrees with cell speeds from micro-channel experiments: (a) Time series of a wild-type mEF cell moving in a microchannel (b) Same as (a) but the images were generated using a computational model. (c) Average cell speed as a function of channel width for the two different cell types.

How is such a trend modified in vimentin-null cells? Since the presence of vimentin makes cell stiffer [71, 72], the vimentin-null cell line is described by a decreased linker spring strength and a decreased nucleus area spring strength, with the latter capturing the lack of a vimentin cage around the nucleus. We indeed observe a similar non-monotonic trend, but with a larger optimal cell speed. Moreover, the optimal cell speed occurs at a narrow channel width, as compared to wild-type cells. Again, as confinement increases, the bulk cytoskeleton in both cell lines also starts deforming (compressing). However, the wild-type cell provides more resistance as it is a stiffer cell, and, thus, pushes against the walls more to effectively act as stronger adhesion to the wall. This effective adhesion to the wall is weaker for the vimentin-null cell line because linker spring strength is weaker. This effect results in both a larger optimal cell speed and the driving force out-competing the adhesion for a larger range of change of channel widths. Note that our results also depend on the nucleus area spring strength. Decreasing the nucleus area spring strength also decreases the effective adhesion to the wall, as the anchoring of the springs to the nucleus is less stiff and so also enhances the cell speed. See Supplementary Fig. S3.

Now let us compare the computational model with the earlier experiments. Our average cell speed is in reasonable quantitative agreement with the experimental cell speed measurements for both cell types and for two different channel widths. See Fig. 2(c). The experiments demonstrated that as confinement increases, vimentin-deficient cell’s average speed also increases, whereas wild-type cell speed remains largely unchanged [32]. We find similar behavior with the non-monotonic trend in cell speed as a function of channel width weaker in the wild-type case. This non-monotonicity also agrees with the experimental observations of other cell lines migrating in the channels [73]. For vimentin-null cells, we predict that for channels less than a 10-micron width that the cell speed will begin to decrease. Hints of this prediction are evident in transwell experiments [25].

### B. Vimentin’s dual role of stress transmitter and nuclear protector

While linker spring strength mediates the competition between driving forces and adhesion forces at the cortex, the linker springs also regulate stress transmission between the actin cortex and the nuclear cortex. One, therefore, may anticipate the shape of the nucleus to depend on the linker spring strength. More specifically, we expect the nucleus to be less responsive to what is happening to the cell cortex so that any shape changes to the nucleus are less dramatic when the linker spring strength is weaker. Prior experiments indeed indicate that the shape of the nucleus is affected by vimentin as cells move in confinement [74, 75]. To test our hypothesis, we compute the shape of the nucleus by finding its circularity C, which is defined as 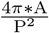, where *A* denotes the cross-sectional area of the nucleus and *P* its cross-sectional perimeter. When C = 1, the nuclear cross-section is a circle. See Fig. 3(a) and (b). As the channel width decreases, there is a greater decrease in C from its essentially surface value for the wild-type cell as compared to the vimentin-null type. This informs us that the wild-type cell, where vimentin is intact, mediates greater stresses to the nucleus as compared to vimentin-null where the cytoskeleton network is weaker, thereby potentially motivating the need for a mesh-like vimentin cage around the nucleus to help mediate the greater stresses.

**FIG. 3.**
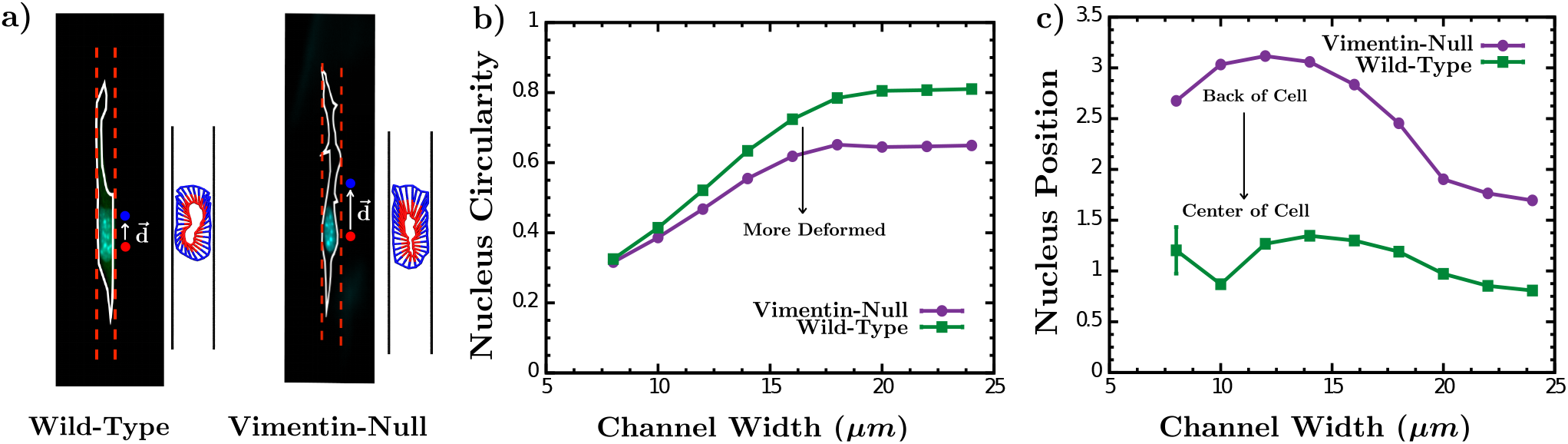
Vimentin affects nuclear shape and position. (a) Images of a wild-type mEF cell and a vimentin-null mEF cell moving in a collagen gel with their computational counterparts. In addition, 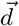 is labeled on each cell image. (b) The circularity of the cell nucleus as a function of confinement for both cell types. (c) The position of the center of mass of the cell nucleus with respect to the center of mass of the cell as a function of channel width for both cell types.

Interestingly, experiments indicate that the nucleus in vimentin-null cells is typically more squishy and has more wrinkles than wild-type, therefore, taking up less effective volume [76]. As discussed in the simulation subsection, in addition to modifying the linker spring strength to go from one cell type to the other, we also modify the area spring constant for the nucleus, with the latter accounting for the mesh-like vimentin cage surrounding the nucleus. When we increase the area spring constant for the nucleus, as well as increase the linker spring stiffness, we find that even though the cytoskeleton still transfers forces from outside to the nucleus with increasing confinement, the nucleus resists such deformations with the circularity tracking very similarly for the smaller channel widths between the two cell types. Moreover, the vimentin-null cells are now always slightly less circular as compared to their wild-type counterpart. The trend is opposite when only modifying the linker spring strength. See Supplementary Fig. S4.

The nuclear envelope is typically viewed as dominated by the nuclear laminas [77, 78]. Here we suggest that vimentin around the nucleus also has a role in stabilizing the shape of the nucleus. Thus, we can conclude that vimentin protects the nucleus from deformations as well as mediates stress transmission between the two cortices, the actin one and the nuclear one, with the latter concept pointed out in Refs. [79, 80]. Now we have quantitative modeling results to substantiate this concept.

In addition to nuclear shape, we computed the center of mass of the nucleus, the center of mass of the cell, and determined the distance between the two, defined as the magnitude of 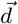 with the origin of 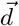 at the center of mass of the nucleus. See Fig. 3(c). This calculation estimates the location of the nucleus with respect to the rest of the cell. When 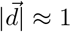, the nucleus is closer to the center of the cell. For larger 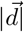, the nucleus moves toward the rear of the cell or away from the leading edge of the cell. We find that the nuclei in the vimentin-null cells are typically positioned toward the rear of the cell, while the nuclei for the wild-type cells are closer to the center of the cell. As the channel width narrows, this difference becomes even starker.

How does this trend of nuclear positioning emerge? For the more malleable linker springs, springs in the leading half of the cell more readily attach to the wall creating a typically flatter leading edge. In doing so, the springs in the leading half of the cell are more extended. From possible range of angles of microtubule polymerization an energetic point-of-view, the extra tension (stretching) in the leading half of the cell is compensated for by the springs in the rear half of the cells configuring to be close to their rest length. Given this energetic argument, one expects this difference in positioning to diminish as temperature increases. This argument also tells us that the nucleus is being pulled by the leading half of the cell as it moves through the channel. We confirm this by calculating the forces on the nucleus due to the cortex. See Supplementary Fig. S4.

### C. The absence of vimentin leads to more polarized cells in nondeformable micro-channels

In addition to cell speed, we also investigate cell direction for both cell types. We propose a novel polarity mechanism based on the position of the nucleus, which is a readout for the position of the centrosome in the cell. The centrosome plays an important role in cell polarity and is also connected to the nucleus via various crosslinkers and proteins [19]. We assume that such connections between the centrosome and nucleus remain strong as the cells move through the channel and so the position of the nucleus tracks the position of the centrosome. We posit in confinement that as the magnitude of 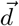 increases, the cell becomes more polarized with the possible range of angles of microtubule polymerization scaling as 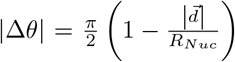, where *R_Nuc_* is the radius of the nucleus and ±Δ*θ* is defined clockwise/counter-clockwise from the reference angle of 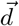 respectively. See Fig. 4(a). There are two reasons for this scaling. First, as the nucleus is more displaced, only the longest microtubules polymerize towards the leading edge of the cell. The lateral confinement and rear end of the cell disrupt microtubule polymerization in the remaining directions. This spatial arrangement biases microtubule polymerization parallel to the confining walls [81–83]. Secondly, vimentin re-organizes itself in about every 15 minutes in wild-type cells. This reorganization can potentially reorient the centrosome. Thus, wild-type cells can potentially alter their direction even in confinement. Given both the position of the nucleus (relative to the rest of the cell) and the presence (or absence) of vimentin, cell repolarization can be very different.

**FIG. 4.**
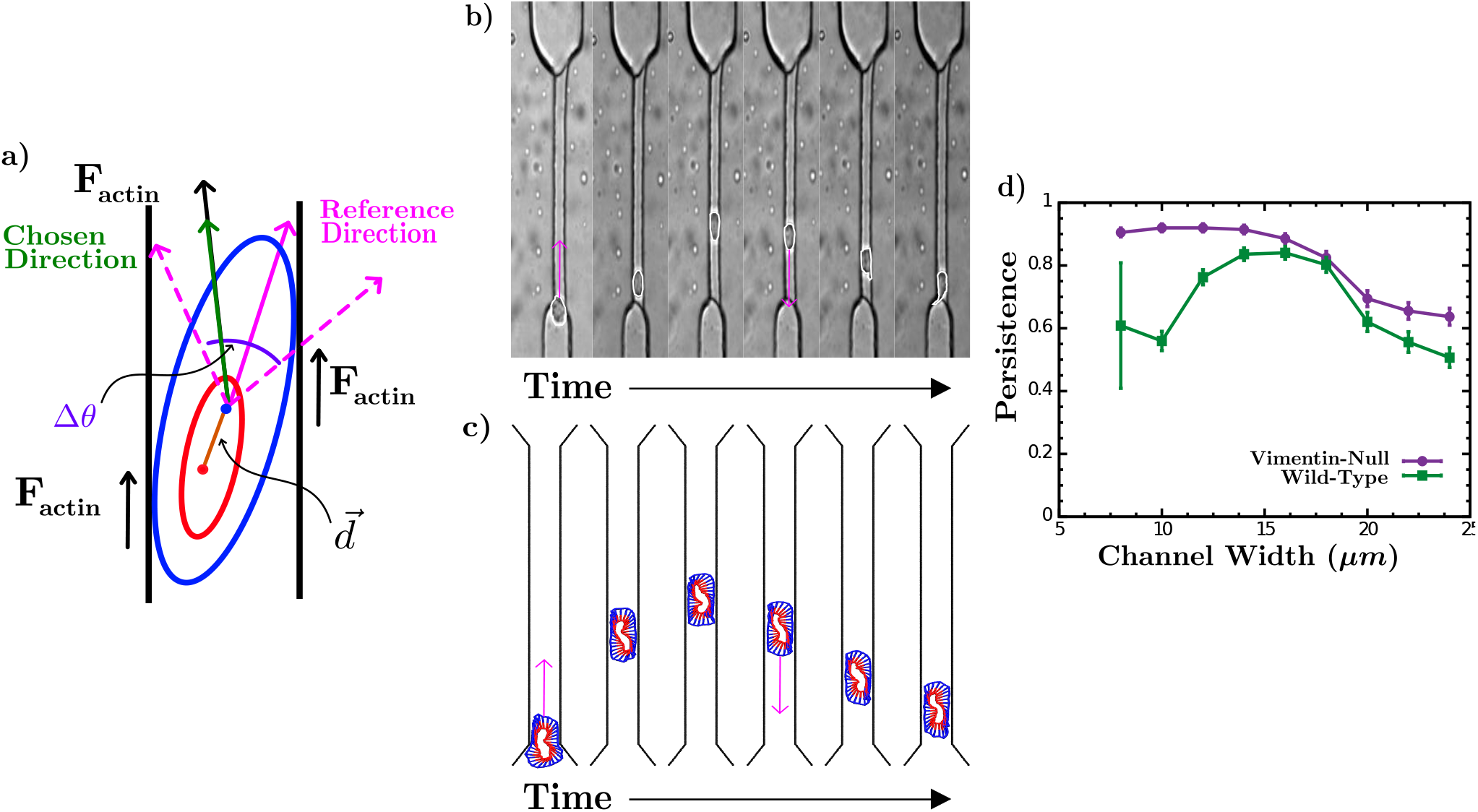
Nucleus-centrosome-based polarity mechanism. (a) Schematic of the polarity mechanism. (b) Time series of a cell going in the channel and changing direction. (c) Computational version of (b). (d) Persistence as a function of channel width for both cell types.

In Figs. 4(b) and (c), we show the time series for a wild-type cell from the earlier micro-channel experiment and compare it with our computational model. As in the experiments, we find that the wild-type cells are more likely to change direction in the channel. To quantify this, we measure the contour length of the trajectory normalized by the length of the micro-channel such that a persistence of unity occurs when the cell does not change direction in the channel. We find that the vimentin-null cell line is more persistent in the channels as compared to the wild-type for all channel widths but more notably different for the smaller channel widths. See. Fig. 4(d). We have also studied the persistence as a function of *k_L_*, 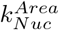, *F_a_*, *F_w_*, and *ϵ_Ad_*. See Supplementary Figs. S5, S6, S7. Given the more asymmetric nuclear positioning in the vimentin-null, a stronger *F_a_* will enhance the tensioning in the leading half of the cell and, thus, decrease Δ*θ* to enhance the polarization, for example.

In the wild-type cells, we note that persistence is initially increasing with increased confinement (Fig. 4(d)). We also see there is a decreased persistence towards the tighter channels. This behavior tracks the non-monotonic behavior observed in the magnitude of 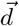 as a function of the channel width. To explain this non-monotonic trend, we turn to the delicate balance between the stiffness of the cell and strain due to channel width. In intermediate channel widths, the wild-type spends less time, thus, have less chance of turning to the other side. The more time wild-type cells spend in the channels, the greater the chance of moving back towards the entrance or getting stuck in the channel. In wider channels, cells do not have enough wall contact but in very narrow channels, stiffness of the cells kicks in and so the cells spend more time in the channel as they travel more slowly in terms of speed.

Indeed, our model recapitulates the experiments. More precisely, experiments found that vimentin-null Fs were more persistent than wild-type mEFs. For example, in 10*μm* channels about half of the wild-type cells did not cross the channel, whereas most of the vimentin-null cells passed to the other side. These results are somewhat surprising as the cells behave opposite on 2D substrates. In 2D, vimentin-deficient cells form lamellipodia in all directions thereby preventing them from polarizing, which is not as likely to occur with their wild-type counterparts.

### D. Vimentin enhances the confinement energy barrier

While experiments do not typically measure energies, we can do so within our model to test the notion of confinement as an energetic barrier to cell motility, particularly since it has been observed that cells tend to migrate in the direction of least confinement to minimize energetic costs [84]. To do so, we compute the time-averaged energy due to conservative potentials while the cell is in the channel. See Fig. 5(a). We repeat this measurement for the different channel widths. See Fig. 5(b). For the two cell lines, the average energy increases more for the wild-type cells than for the vimentin-null cells as the confinement increases, as anticipated. However, the increase in energy is non-linear. Interestingly, recent modeling studies the phenomenon of compression stiffening in cells. Compression stiffening, a nonlinear rheological property in which a material’s moduli increase with increasing uniaxial compressive strain, has recently been discovered in static cells [85]. Such a phenomenon should also manifest itself in confined cell motility. We have measured the compressive strain in the cells and find a good correlation between compressive strain and channel width (see Supplementary Fig. S9). Since the derivative of the energy with respect to strain, or channel width, relates to the compressive stress of the cell, should a quantity proportional to the compressive stress increase faster than linear with the decreasing channel width, then the motile cells are indeed exhibiting compression stiffening. We find this to be the case. See. Fig. 5(c). Since the vimentin-null are more deformable, their compression stiffen is less dramatic and the onset occurs at a slightly higher strain.

**FIG. 5.**
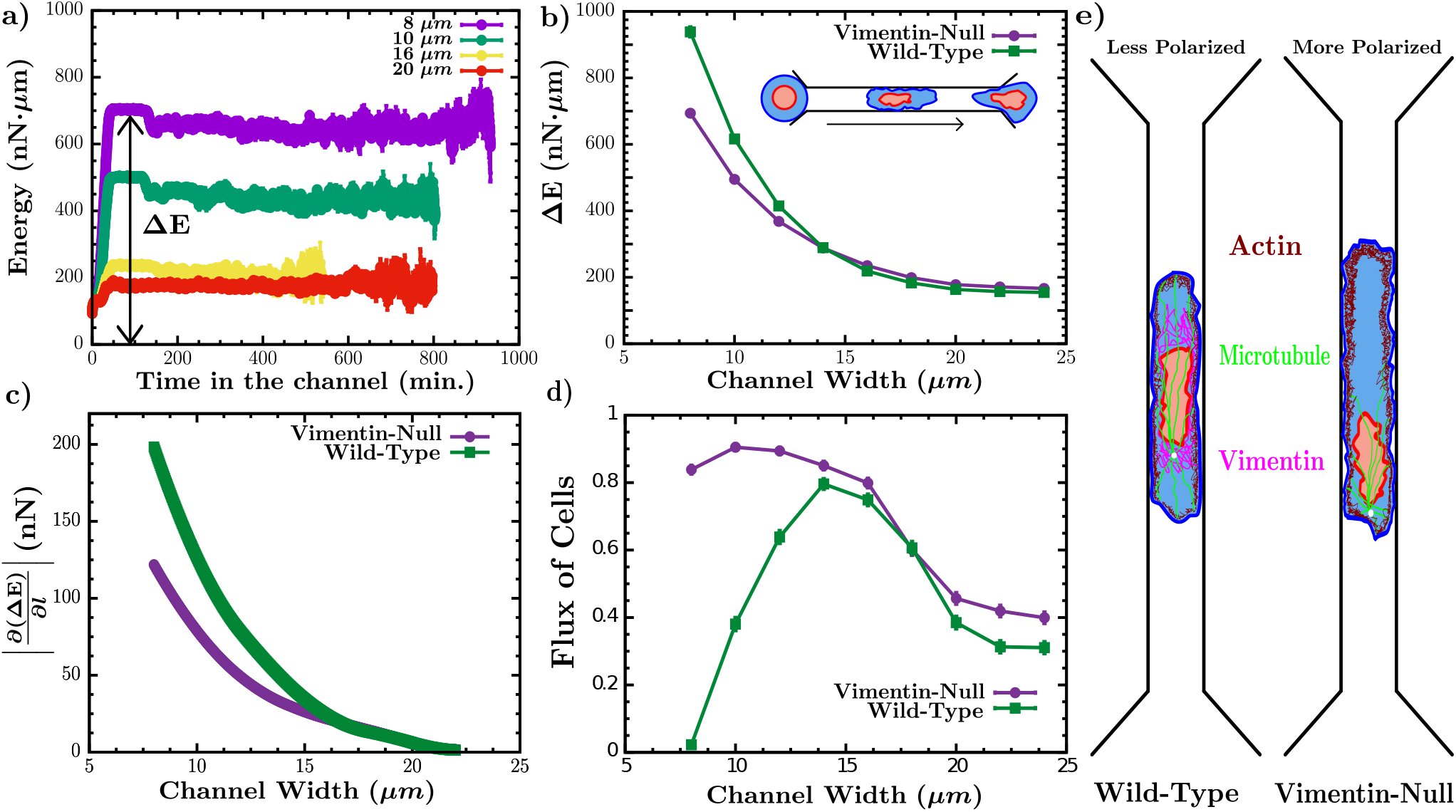
Energy barriers in confined cell motility. (a) The average energy of each cell due to the conservative forces as a function of time for different channel widths. (b) The time-averaged energy as a function of channel width for both cell types. (c) The magnitude of the numerical derivative of the time-averaged energy as a function of channel width for both cell types. (d) The flux of each cell type as a function of the channel width. (e) A schematic of the internal organization of more polarized/persistent cells versus less polarized/persistent ones.

In addition to the energetic barrier, there is also a time scale for entering (versus crossing) the channel translates into a rate for attempting to hop over the energy barrier. This attempt rate depends on the polarization of the cell. Prior to entering the channel, the cell effectively sees a two-dimensional surface. In this case, the magnitude of 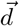 is smaller (for both cell types), so that we anticipate more changes in cell direction. Since we initialize the cells to move towards the channel opening, we do not explore the attempt rate here. In going to 2D, we anticipate that the polarizability of the cell changes from our new mechanism to one that depends on the fluctuations in *d*, as opposed to the average. Moreover, we anticipate the fluctuations in *d* being larger in the vimentin-null case, which corresponds to more possible directions in the cells.

While we do not explore the attempt rate here, the non-linear increase in the energy barrier as confinement increases accounts for the dramatic drop in the flux, or the fraction of the cells exiting out the other side, as a function of channel width for the wild-type cells. For those narrow channels, the wild-type cells are not able to enter the channel in the first place while for the vimentin-null, they are, and hence the dramatic drop in flux for the smaller channel widths. See Fig. 5(d). Such behavior agrees with experimental observations.

## IV. DISCUSSION

In focusing on vimentin’s role in confined cell motility using a computational model that captures the roles of *all three types of cytoskeletal filaments as well as the nucleus*, we investigate the dual role vimentin plays, the first being the mechanical protector of the nucleus and the second being a stress regulator between the outer and inner cortices. For the first role, we modify the stiffness of the nucleus, and for the second, we modify the stiffness of the mechanical connections, or linker springs, between the inner and outer cortices. More precisely, in going from wild-type to vimentin-null, we weaken both stiffnesses. For the wild-type cells, we find a nonmonotonic dependence of cell speed with channel width. As the channel width narrows, the cell’s cortex increases its contact with the wall, which, in turn, generates more driving force to increase cell speed. Yet, this trend competes with the bulk deformability of the cell via the linker springs and nucleus to increase the effective adhesion as the channel width decreases, leading to a slower cell speed. For the vimentin-null cells, we observe a similar non-monotonic trend, but with a larger optimal cell speed. Moreover, the optimal cell speed occurs at a narrow channel width, as compared to wild-type cells, given the enhanced bulk deformability of the cells. Thus, for nondeformable confinement with simple geometry, vimentin-null cell speed is typically faster, which is seemingly contrary to the notion that the EMT cells typically upregulate vimentin to be able to move more efficiently.

And yet, there is another ingredient to cell motility beyond the speed, which is cell direction, or cell polarity. Given our proposition that the larger the distance between the center of mass of the cell and the center of mass of the nucleus, the more polarized, or directed, a cell is, we determine that vimentin-null mEFs are more polarized than wild-type cells. This trend emerges because the nucleus is typically located more to the rear of the cell in the vimentin-null case with the nucleus being pulled by the leading edge of the cell as it travels through the microchannel. The enhanced polarization indicates that the vimentin-null cells are more subjugated to the confinement since their own internal polarization mechanism that depends on cross-talk of the centrosome with the nucleus and other cytoskeletal connections are diminished. In other words, the *vimentin-null cells rely more heavily on external cues*, at least in this stiff microchannel environment. See Fig. 5(e). We also find typically different nuclear shapes between the two cases for wider channel widths with the vimentin-null cell nuclei less circular. As the channel widths become increasingly smaller, the circularity of each nucleus type becomes more similar. Finally, since energetic costs are known to be a predictor of the migration path in confined cell motility, we find a higher nonlinear energy barrier for wild-type cells entering more confined channels as compared to the vimentin-null cells, which, again, is seemingly contrary to the notion of the upregulation of vimentin enhancing cell motility.

What do our findings tell us about the interaction between a cell and its microenvironment more generally? In the absence of vimentin, the cells become more deformable and so more mechanically sensitive to their microenvironment. In the event that the cell’s microenvironment is very much like a microchannel with a simple linear geometry, then downregulation of vimentin helps the cell travel more effectively from one place to another in the microenvironment. However, in a more generic microenvironment, such a setup is not as likely. There-fore, for a cell to upregulate vimentin in order to enhance motility, which is the usual canon, translates to a cell that is enhancing its own internal polarization mechanism to effectively search the microenvironment for a minimal energy barrier by being able to more readily able to change direction. In other words, the cell is less enslaved to the “whims” of the microenvironment. Cells can do this despite the increase in the energy barrier because they have developed coping mechanisms, such as Arp2/3 branched actin near the cell nucleus helping it to squeeze through small pores [86], which are not currently accounted for in the model. Additionally, with the upregulation of vimentin, there is more mechanical cross-talk between the two cortices to perhaps increase the role of the nucleus itself in regulating cell mechanics. Recent work suggests that compressed nuclei release calcium into the cytoplasm to help reconfigure the cytoskeleton [87, 88]. Therefore, our theoretical findings provide a much richer interpretation of how vimentin affects cell migration with the combination of stress coupler between inner and outer parts of the cell, nuclear protector, and now polarity regulator. Perhaps keratin is typically downregulated in the EMT because it is not able to simultaneously perform all three roles.

How does our model compare with other models of confined cell motility? Active gel models of the cells predict that cells repel their way through confined spaces much like a climber repelling off a wall [89]. Such a model does not explicitly include a nucleus nor a centrosome such that cell polarity is an input. Another model based on the molecular-clutch mechanism explores glial cells moving through microchannels [90]. Cell polarity is, again, an input. A third model with the complexity of an actomyosin cortex and a nucleus and couplings in between demonstrates that there is a nonmonotonic relationship between cell speed and matrix stiffness in two-dimensions [91]. Cell confinement in this model is explored by constraining the beads/nodes to stay within a channel geometry. The cell direction of motion is randomly chosen every so many minutes. While cell speed is studied, cell persistence is not. Here, cell polarization emerges from an intra-cellular detail rooted in the position of the centrosome assumed to be in close proximity with the nucleus. There are indeed additional models, notably, Refs. [92, 93]. Our model walks the fine line between being minimal and yet detailed enough to quantify the new functionality of vimentin.

While we have focused on comparison with mEFs, we expect our findings to generalize to other mesenchymal cell types given our generic model. Specifically, another cell type would presumably translate to a slightly different set of parameters. Since our findings are robust for a range of several parameters, we expect our conclusions to generalize to at least some cell types. Given the paradigm of the EMT, we aim to test our predictions on cell types that do undergo an EMT transition while also accounting for the typical downregulation of keratin. Since we have focused here on mesenchymal cells, it would be interesting to generalize our model to non-mesenchymal cells and explore the role of vimentin in other motility modes of migration [94]. It would also be interesting to alter the geometry of the microchannel as well as study multiple cells moving in confinement to determine the robustness of our findings. Extending our polarization mechanism to include multicellular interactions will shed more light on the phenomenon of contact inhibition of locomotion in which motile cells stop moving or change direction upon contact with another cell. Furthermore, it has been also found that matrix stiffness has a part to play in positioning MTOC [95] which would also be intriguing to explore.

## V. ACKNOWLEDGEMENTS

JMS acknowledges NSF-DMR-1832002 and an Isaac Newton Award from the DoD. JMS and AEP acknowledge a Syracuse University CUSE grant and a Syracuse BioInspired grant. SG acknowledges a graduate fellowship from Syracuse University.

## VI. SUPPLEMENTARY FIGURES

**FIG. S1.**
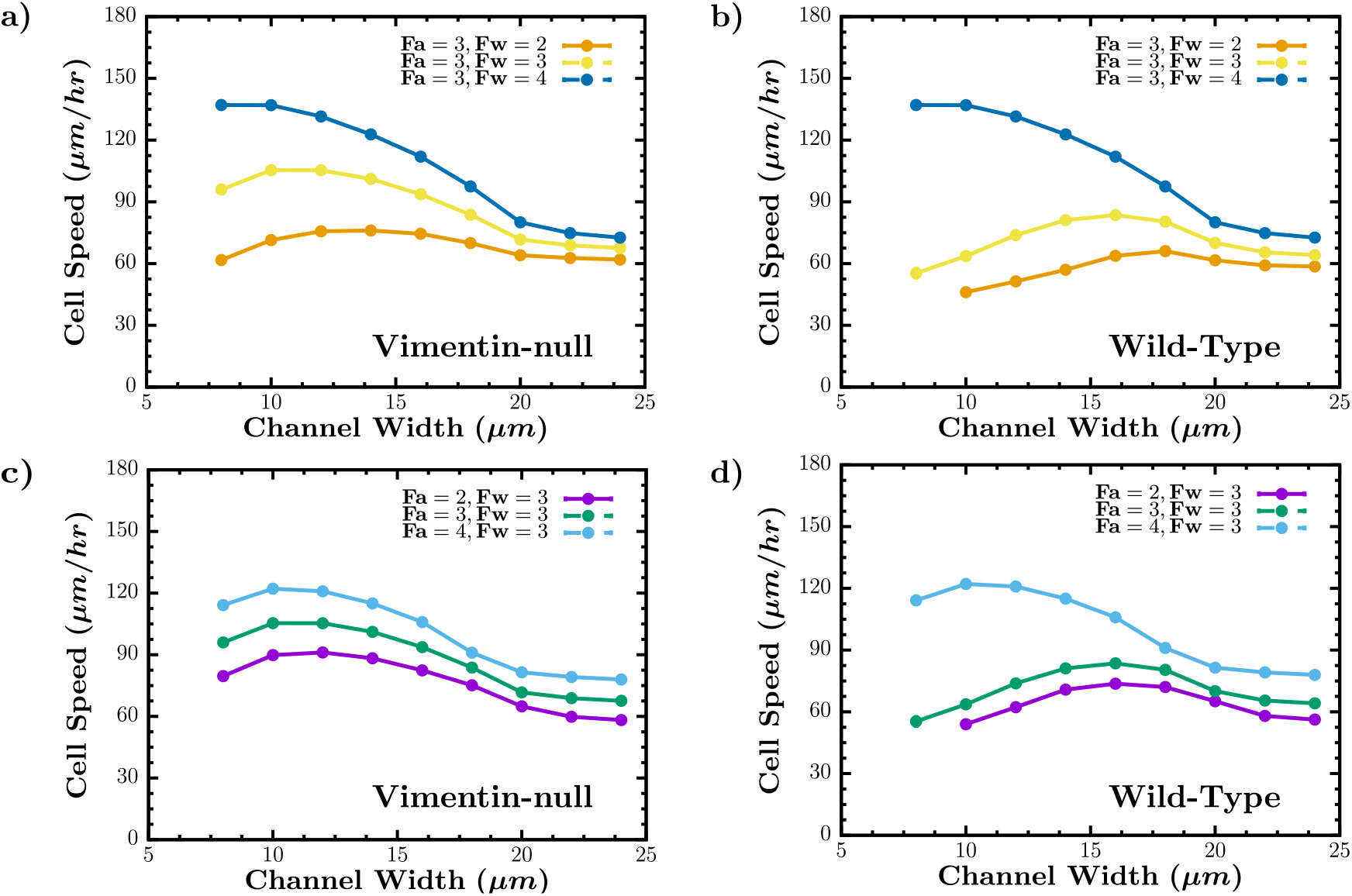
Cell speed as a function of channel width for different actin forces at the leading edge and actin forces at the wall.’ In (a) and (b), the magnitude of the actin force at the leading edge *F_a_* is fixed and the magnitude of the actin force at the wall *F_w_* is varied for each cell type and as a function of channel width. In (c) and (d), the magnitude of the actin force at the leading edge is now varied.

**FIG. S2.**
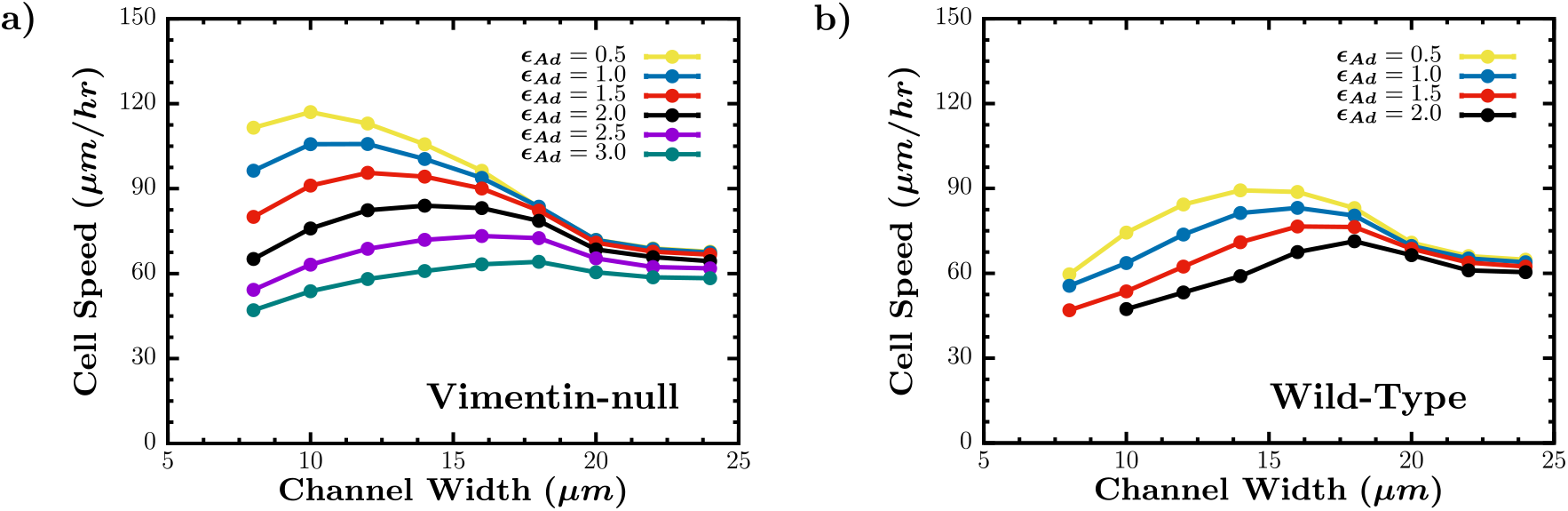
Cell speed as a function of channel width for different adhesion strengths. (a) Vimentin-null cell (b) Wild-type cell.

**FIG. S3.**
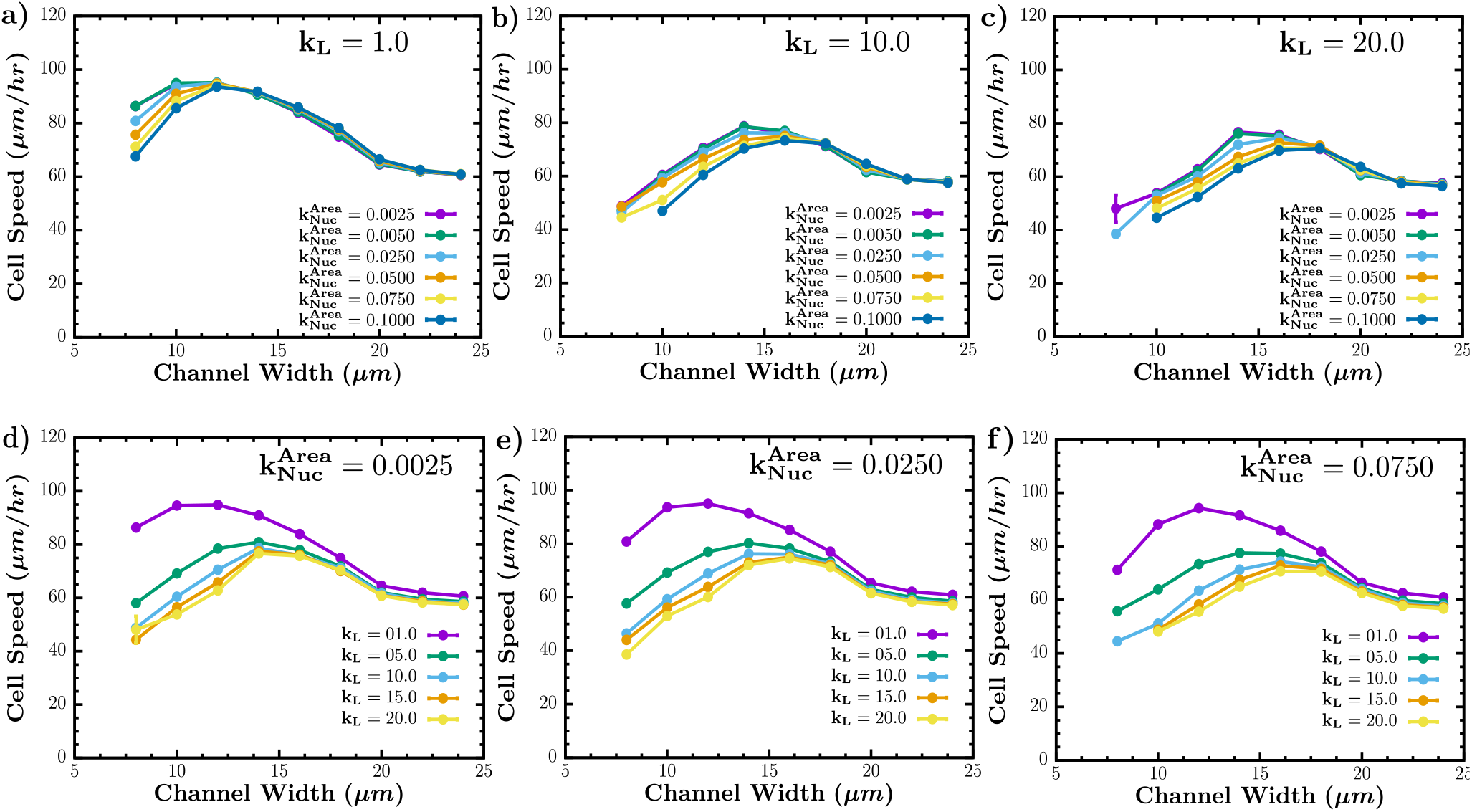
Cell Speed as a function of channel width for different linker and nuclear area spring strengths: (a)-(c) Varying nuclear area spring strength 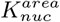 for different linker spring strengths, *K_L_*. (d)-(f) Varying linker spring strengths, *K_L_*, for different nuclear area spring strengths 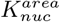.

**FIG. S4.**
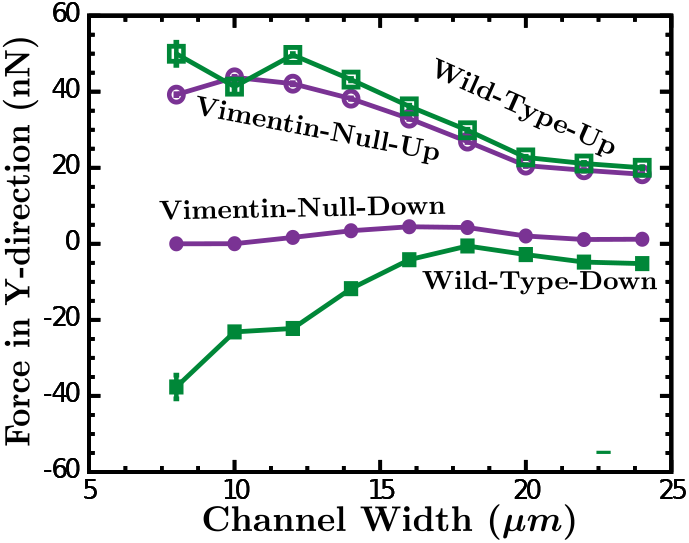
Forces on the nucleus due to the actomyosin cortex: The net force in the y-direction on the top (upper) half of the nucleus due to the leading half of the actomyosin cortex and the net force in the y-direction on the bottom (lower) half of the nucleus due to the rear half of the actomyosin cortex as a function of channel width for each cell type.

**FIG. S5.**
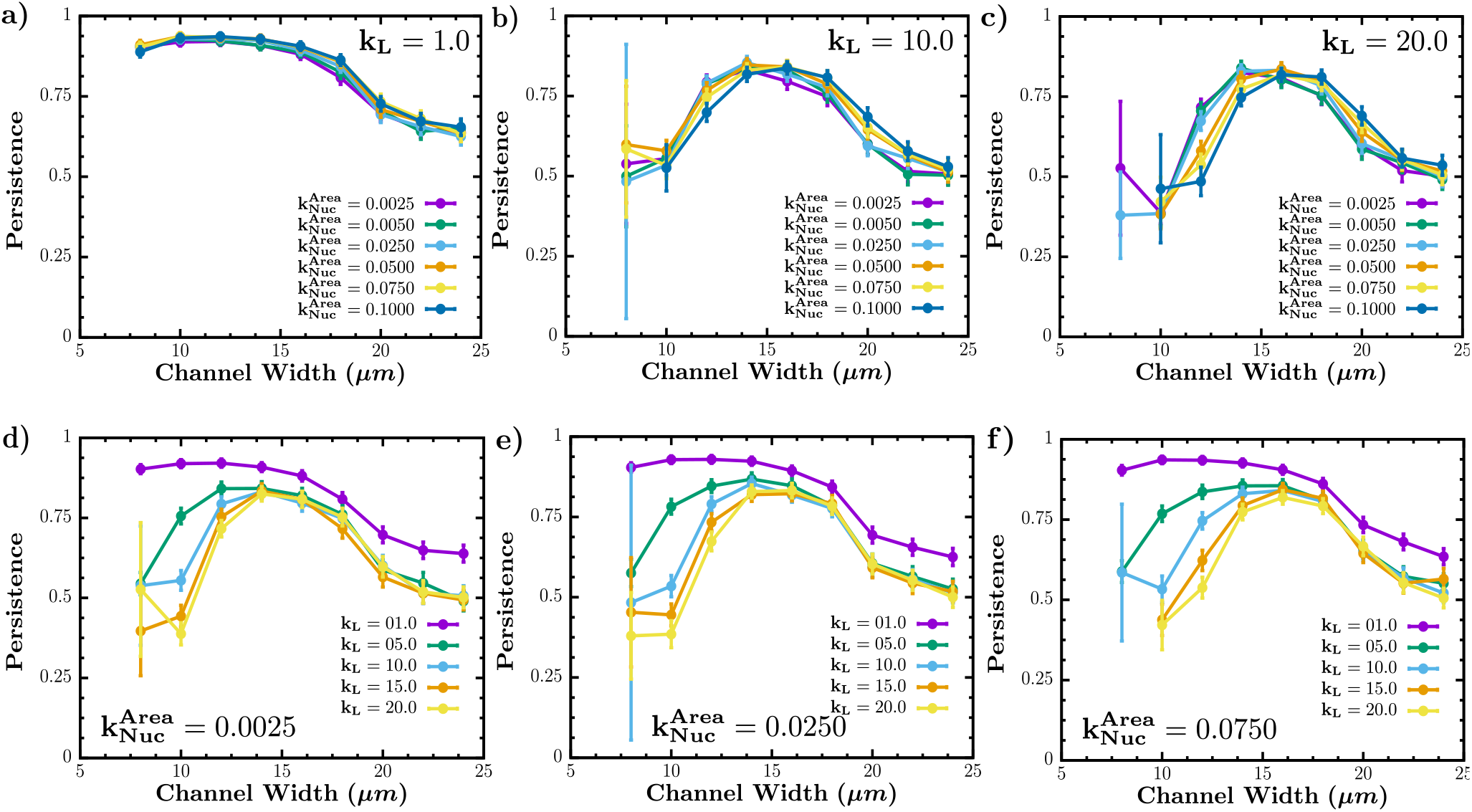
Persistence as a function of channel width for different linker and nuclear area spring strengths: (a)-(c) Varying nuclear area spring strength 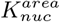 for different linker spring strengths, *K_L_*. (d)-(f) Varying linker spring strengths, *K_L_*, for different nuclear area spring strengths 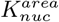.

**FIG. S6.**
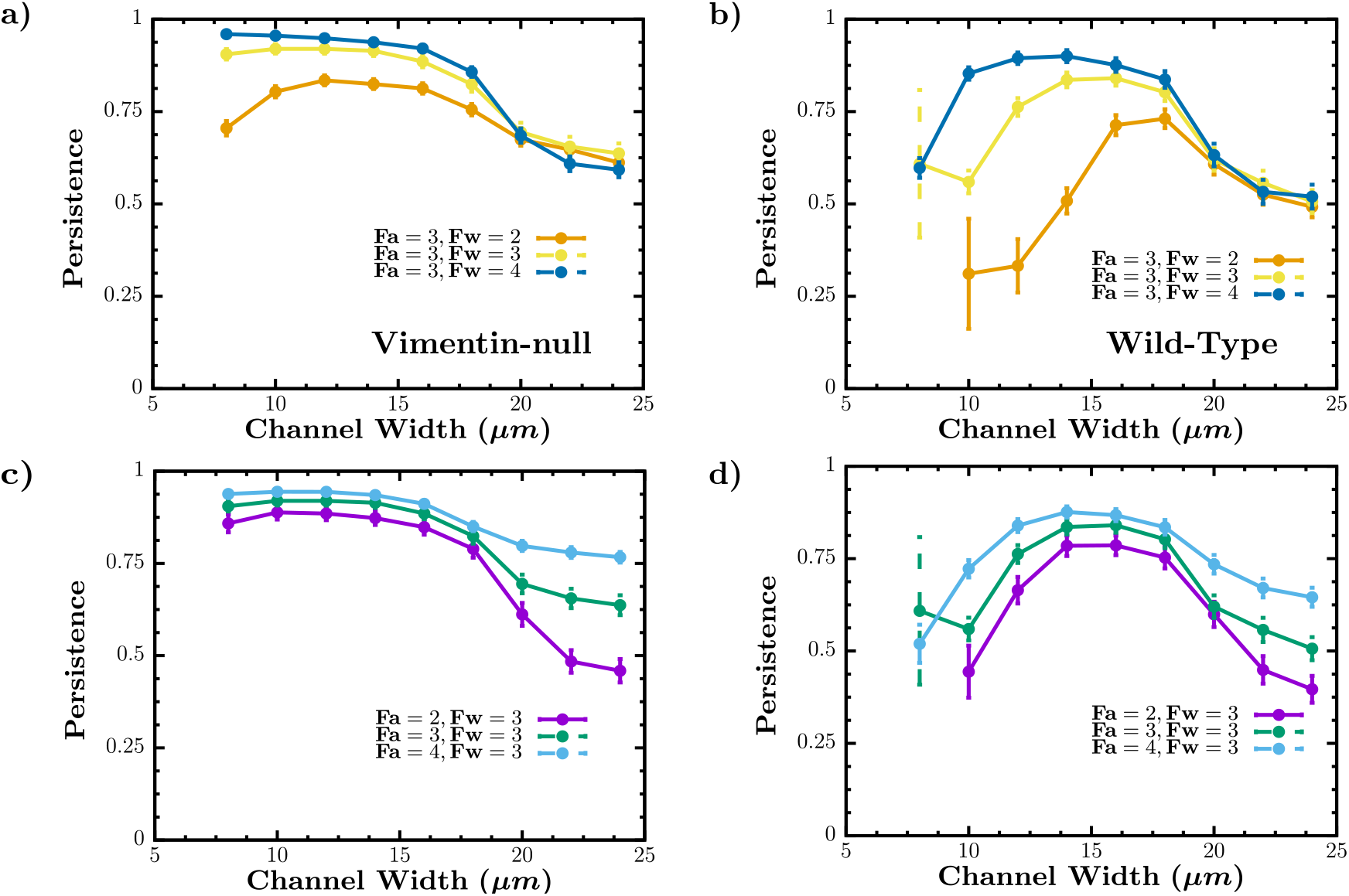
Persistence as a function of channel width for different actin forces at the leading edge and actin forces at the wall.’ In (a) and (b), the magnitude of the actin force at the leading edge *F_a_* is fixed and the magnitude of the actin force at the wall *F_w_* is varied for each cell type and as a function of channel width. In (c) and (d), the magnitude of the actin force at the leading edge is now varied.

**FIG. S7.**
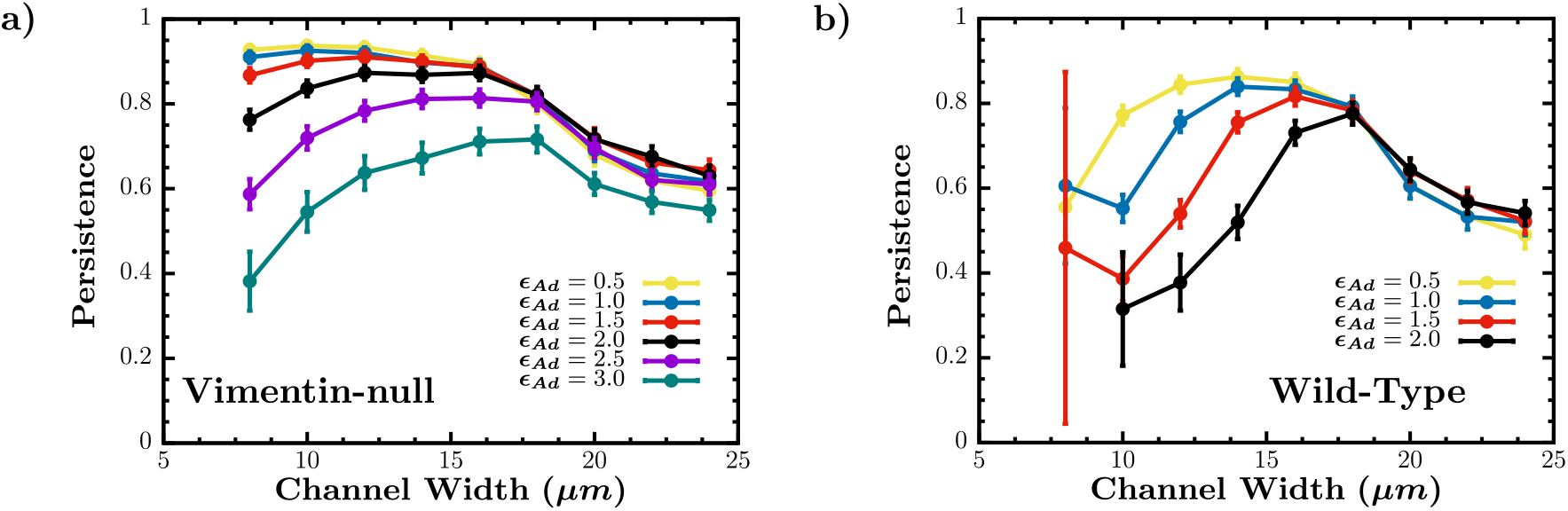
Persistence as a function of channel width for different adhesion strengths. (a) Vimentin-null cell (b) Wild-type cell.

**FIG. S8.**
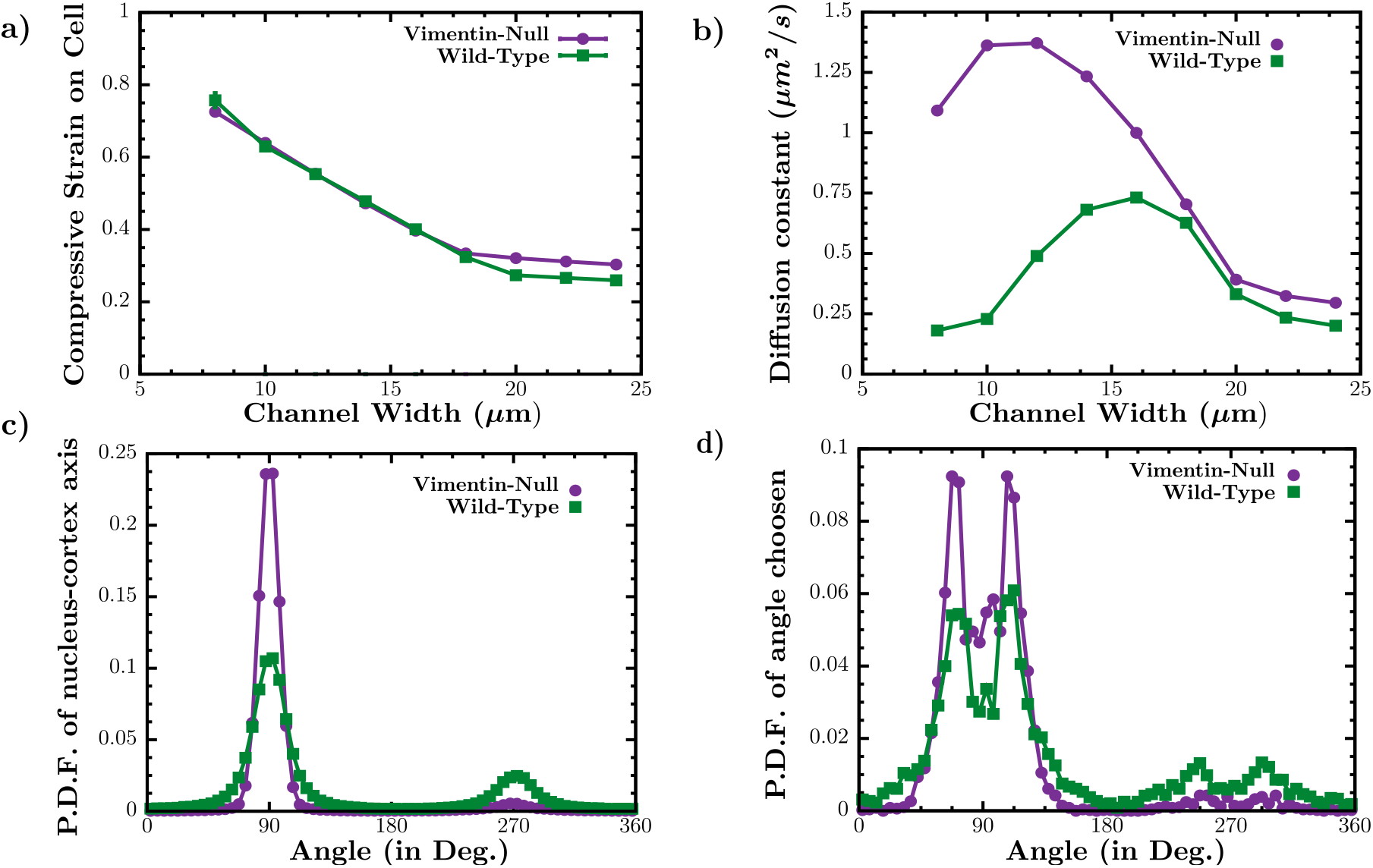
Additional properties. (a) Compressive strain on the cell versus channel width for both cell types. (b) Translational diffusion constant for both cell lines. (c) Probability distribution function of the angle between x-axis and the Nucleus-axis for both cell types. (d) Probability distribution function of the angle choosen by the cell (with respect to the x-axis) for a 10 *μ*m channel width.

## References

[1] C. Leduc and S. Etienne-Manneville, Current Opinion in Cell Biology 32, 102 (2015).

[2] F. Danielsson, M. Peterson, H. Caldeira Araújo, F. Laut-enschläger, and A. Gad, Cells 7, 147 (2018).

[3] M. Pekny and E. B. Lane, Experimental Cell Research 313, 2244 (2007).

[4] B. T. Helfand, M. G. Mendez, S. N. Murthy, D. K. Shumaker, B. Grin, S. Mahammad, U. Aebi, T. Wedig, Y. I. Wu, K. M. Hahn, M. Inagaki, H. Herrmann, and R. D. Goldman, Molecular Biology of the Cell 22, 1274 (2011).

[5] M. G. Mendez, S. Kojima, and R. D. Goldman, The FASEB Journal 24, 1838 (2010).

[6] B. Eckes, E. Colucci-Guyon, H. Smola, S. Nodder, C. Babinet, T. Krieg, and P. Martin, Journal of Cell Science 113, 2455 (2000).

[7] M. R. Rogel, P. N. Soni, J. R. Troken, A. Sitikov, H. E. Trejo, and K. M. Ridge, The FASEB Journal 25, 3873 (2011).

[8] M. E. Kidd, D. K. Shumaker, and K. M. Ridge, American Journal of Respiratory Cell and Molecular Biology 50, 1 (2014).

[9] A. Satelli and S. Li, Cellular and Molecular Life Sciences 68, 3033 (2011).

[10] G. Danuser, J. Allard, and A. Mogilner, Annual Review of Cell and Developmental Biology 29, 501 (2013).

[11] B. Winkler, I. S. Aranson, and F. Ziebert, Communications Physics 2(2019), 10.1038/s42005-019-0185-x.

[12] S. Hervas-Raluy, J. M. Garcia-Aznar, and M. J. Gomez-Benito, Biomechanics and Modeling in Mechanobiology 18, 1177 (2019).

[13] C. Abaurrea-Velasco, T. Auth, and G. Gompper, arXiv (2018), arXiv:1812.09932.

[14] Y. Cao, R. Karmakar, E. Ghabache, E. Gutierrez, Y. Zhao, A. Groisman, H. Levine, B. A. Camley, and W. J. Rappel, Soft Matter 15, 2043 (2019), arXiv:1806.01315.

[15] M. Wang, B. Cheng, Y. Yang, H. Liu, G. Huang, L. Han, F. Li, and F. Xu, Nano Letters 19, 5949 (2019).

[16] B. P. Bouchet and A. Akhmanova, Journal of Cell Science 130, 39 (2017).

[17] Z. L. Zhao, Z. Y. Liu, J. Du, G. K. Xu, and X. Q. Feng, Biophysical Journal 112, 2377 (2017).

[18] D. M. Graham, T. Andersen, L. Sharek, G. Uzer, K. Rothenberg, B. D. Hoffman, J. Rubin, M. Balland, J. E. Bear, and K. Burridge, Journal of Cell Biology 217, 895 (2018).

[19] G. W. Luxton and G. G. Gundersen, Current Opinion in Cell Biology 23, 579 (2011).

[20] P. Friedl and E. B. Brocker, “The biology of cell locomotion within three-dimensional extracellular matrix,” (2000).

[21] R. J. Petrie and K. M. Yamada, Journal of Cell Science 125, 5917 (2012).

[22] E. M. Balzer, Z. Tong, C. D. Paul, W. C. Hung, K. M. Stroka, A. E. Boggs, S. S. Martin, and K. Konstantopoulos, FASEB Journal 26, 4045 (2012).

[23] A. D. Doyle, F. W. Wang, K. Matsumoto, and K. M. Yamada, Journal of Cell Biology 184, 481 (2009).

[24] J. Zhang and Y.-l. Wang, Molecular Biology of the Cell 28, 3240 (2017).

[25] A. E. Patteson, A. Vahabikashi, K. Pogoda, S. A. Adam, K. Mandal, M. Kittisopikul, S. Sivagurunathan, A. Goldman, R. D. Goldman, and P. A. Janmey, The Journal of cell biology 218, 4079 (2019).

[26] A. Reversat, F. Gaertner, J. Merrin, J. Stopp, S. Tasciyan, J. Aguilera, I. de Vries, R. Hauschild, M. Hons, M. Piel, A. Callan-Jones, R. Voituriez, and M. Sixt, Nature 582, 582 (2020).

[27] A. Moure and H. Gomez, Biomechanics and Modeling in Mechanobiology 19, 1491 (2020).

[28] D. Aubry, H. Thiam, M. Piel, and R. Allena, Biomechanics and Modeling in Mechanobiology 14, 143 (2015).

[29] C. Giverso, A. Grillo, and L. Preziosi, Biomechanics and Modeling in Mechanobiology 13, 481 (2014).

[30] M. Le Berre, Y. J. Liu, J. Hu, P. Maiuri, O. Bénichou, R. Voituriez, Y. Chen, and M. Piel, Physical Review Letters 111, 1 (2013), arXiv:1310.4129.

[31] N. Costigliola, L. Ding, C. J. Burckhardt, S. J. Han, E. Gutierrez, A. Mota, A. Groisman, T. J. Mitchison, and G. Danuser, Proceedings of the National Academy of Sciences of the United States of America (2017), 10.1073/pnas.1614610114.

[32] A. E. Patteson, K. Pogoda, F. J. Byfield, K. Mandal, Z. Ostrowska-Podhorodecka, E. E. Charrier, P. A. Galie, P. Deptula, R. Bucki, C. A. McCulloch, and P. A. Janmey, Small 15, 1 (2019).

[33] R. A. Battaglia, S. Delic, H. Herrmann, and N. T. Snider, F1000Research 7, 1796 (2018).

[34] L. D. C. Stankevicins, M. Urbanska, D. A. Flormann, E. Terriac, Z. Mostajeran, A. Gad, F. Cheng, J. Eriksson, and F. Lautenschläger, 1 (2019).

[35] E. J. van Bodegraven and S. Etienne-Manneville, Current Opinion in Cell Biology 66, 79 (2020).

[36] Y. Jiu, J. Peränen, N. Schaible, F. Cheng, J. E. Eriksson, R. Krishnan, and P. Lappalainen, Journal of Cell Science 130, 892 (2017).

[37] Z. Gan, L. Ding, C. J. Burckhardt, J. Lowery, A. Zaritsky, K. Sitterley, A. Mota, N. Costigliola, C. G. Starker, D. F. Voytas, J. Tytell, R. D. Goldman, and G. Danuser, Cell Systems 3, 500 (2016).

[38] A. E. Patteson, A. Vahabikashi, R. D. Goldman, and P. A. Janmey, BioEssays 2000078, 1 (2020).

[39] A. E. Patteson, R. J. Carroll, D. V. Iwamoto, and P. A. Janmey, Physical Biology 18 (2021), 10.1088/1478-3975/abbcc2.

[40] J. P. Thiery, H. Acloque, R. Y. Huang, and M. A. Nieto, Cell 139, 871 (2009).

[41] P. A. Janmey, U. Euteneuer, P. Traub, and M. Schliwa, Journal of Cell Biology 113, 155 (1991).

[42] M. Guo, A. J. Ehrlicher, S. Mahammad, H. Fabich, M. H. Jensen, J. R. Moore, J. J. Fredberg, R. D. Goldman, and D. A. Weitz, Biophysical Journal 105, 1562 (2013).

[43] A. A. Minin and M. V. Moldaver, Biochemistry (Moscow) 73, 1453 (2008).

[44] L. Chang, K. Barlan, Y. H. Chou, B. Grin, M. Lakonishok, A. S. Serpinskaya, D. K. Shumaker, H. Herrmann, V. I. Gelfand, and R. D. Goldman, Journal of Cell Science 122, 2914 (2009).

[45] P. M. Davidson, C. Denais, M. C. Bakshi, and J. Lammerding, Cellular and Molecular Bioengineering 7, 293 (2014).

[46] S. Seetharaman and S. Etienne-Manneville, Trends in Cell Biology 30, 720 (2020).

[47] O. Esue, A. A. Carson, Y. Tseng, and D. Wirtz, Journal of Biological Chemistry 281, 30393 (2006).

[48] T. M. Svitkina, A. B. Verkhovsky, and G. B. Borisy, Biological Bulletin 194, 409 (1998).

[49] S. Osmanagic-Myers, S. Rus, M. Wolfram, D. Brunner, W. H. Goldmann, N. Bonakdar, I. Fischer, S. Reipert, A. Zuzuarregui, G. Walko, and G. Wiche, Journal of Cell Science 128, 4138 (2015).

[50] J. Liu, T. R. Ben-Shahar, D. Riemer, M. Treinin, P. Spann, K. Weber, A. Fire, and Y. Gruenbaum, Molecular Biology of the Cell 11, 3937 (2000).

[51] J. Lammerding, P. C. Schulze, T. Takahashi, S. Kozlov, T. Sullivan, R. D. Kamm, C. L. Stewart, and R. T. Lee, Journal of Clinical Investigation 113, 370 (2004).

[52] M. Crisp, Q. Liu, K. Roux, J. B. Rattner, C. Shanahan, B. Burke, P. D. Stahl, and D. Hodzic, Journal of Cell Biology 172, 41 (2006).

[53] K. J. Roux, M. L. Crisp, Q. Liu, D. Kim, S. Kozlov, C. L. Stewart, and B. Burke, Proceedings of the National Academy of Sciences 106, 2194 (2009).

[54] K. Wilhelmsen, S. H. M. Litjens, I. Kuikman, N. Tshimbalanga, H. Janssen, I. Van Den Bout, K. Raymond, A. Sonnenberg, I. D. Van Bout, K. Raymond, and A. Sonnenberg, Journal of Cell Biology 171, 799 (2005).

[55] M. L. Lombardi, D. E. Jaalouk, C. M. Shanahan, B. Burke, K. J. Roux, and J. Lammerding, Journal of Biological Chemistry 286, 26743 (2011).

[56] S. B. Khatau, R. J. Bloom, S. Bajpai, D. Razafsky, S. Zang, A. Giri, P. H. Wu, J. Marchand, A. Celedon, C. M. Hale, S. X. Sun, D. Hodzic, and D. Wirtz, Scientific Reports 2 (2012), 10.1038/srep00488.

[57] B. L. Bangasser and D. J. Odde, Cellular and Molecular Bioengineering 6, 449 (2013).

[58] B. L. Bangasser, S. S. Rosenfeld, and D. J. Odde, Biophysical Journal 105, 581 (2013).

[59] C. E. Chan and D. J. Odde, Science 322, 1687 (2008).

[60] E. A. E. Calderwood and D. A., 316, 1148 (2007).

[61] J. C. Meiring, B. I. Shneyer, and A. Akhmanova, Current Opinion in Cell Biology 62, 86 (2020).

[62] C. D. Paul, W.-C. Hung, D. Wirtz, and K. Konstantopoulos, Annual Review of Biomedical Engineering 18, 159 (2016).

[63] S. H. Shabbir, M. M. Cleland, R. D. Goldman, and M. Mrksich, Biomaterials 35, 1359 (2014).

[64] N. Caille, O. Thoumine, Y. Tardy, and J. J. Meister, Journal of Biomechanics 35, 177 (2002).

[65] A. Vahabikashi, Biophysical Journal 91, 5753 (2019).

[66] H. V. Prentice-Mott, C. H. Chang, L. Mahadevan, T. J. Mitchison, D. Irimia, and J. V. Shah, Proceedings of the National Academy of Sciences of the United States of America 110, 21006 (2013).

[67] N. M. Wakida, E. L. Botvinick, J. Lin, and M. W. Berns, PLoS ONE 5, e15462 (2010).

[68] A. D. Doyle, F. W. Wang, K. Matsumoto, and K. M. Yamada, Journal of Cell Biology 184, 481 (2009).

[69] M. Ueda, R. Gräf, H. K. Macwilliams, M. Schliwa, and U. Euteneuer, Proceedings of the National Academy of Sciences of the United States of America 94, 9674 (1997).

[70] G. Salpingidou, A. Smertenko, I. Hausmanowa-Petrucewicz, P. J. Hussey, and C. J. Hutchison, The Journal of cell biology 178, 897 (2007).

[71] B. Eckes, D. Dogic, E. Colucci-Guyon, N. Wang, A. Maniotis, D. Ingber, A. Merckling, F. Langa, M. Aumailley, A. Delouvée, V. Koteliansky, C. Babinet, and T. Krieg, Journal of cell science 111 ( Pt 1, 1897 (1998).

[72] N. Wang and D. Stamenovic, Journal of Muscle Research and Cell Motility 23, 535 (2002).

[73] G. Charras and E. Sahai, Nature Reviews Molecular Cell Biology 15, 813 (2014).

[74] I. Dupin, Y. Sakamoto, S. Etienne-Manneville, A. J. Sar-ria, J. G. Lieber, S. K. Nordeen, and R. M. Evans, Journal of Cell Science 107, 1593 (1994).

[75] J. Lowery, E. R. Kuczmarski, H. Herrmann, and R. D. Goldma, Journal of Biological Chemistry 290, 17145 (2015).

[76] T. P. Lele, R. B. Dickinson, and G. G. Gundersen, Journal of Cell Biology 217, 3330 (2018).

[77] A. L. McGregor, C. R. Hsia, and J. Lammerding, Current Opinion in Cell Biology 40, 32 (2016).

[78] J. Renkawitz and M. Sixt, Cell 167, 1448 (2016).

[79] A. J. Maniotis, C. S. Chen, and D. E. Ingber, Proceedings of the National Academy of Sciences of the United States of America 94, 849 (1997).

[80] S. Neelam, T. J. Chancellor, Y. Li, J. A. Nickerson, K. J. Roux, R. B. Dickinson, and T. P. Lele, Proceedings of the National Academy of Sciences of the United States of America 112, 5720 (2015).

[81] M. C. Lagomarsino, C. Tanase, J. W. Vos, A. M. C. Emons, B. M. Mulder, and M. Dogterom, Biophysical Journal 92, 1046 (2007).

[82] M. Pinot, F. Chesnel, J. Z. Kubiak, I. Arnal, F. J. Nedelec, and Z. Gueroui, Current Biology 19, 954 (2009).

[83] P. B. Desai, J. R. Freshour, D. R. Mitchell, and C. Author, , 1 (2014).

[84] M. R. Zanotelli, A. Rahman-Zaman, J. A. VanderBurgh, P. V. Taufalele, A. Jain, D. Erickson, F. Bordeleau, and C. A. Reinhart-King, Nature Communications 10, 1 (2019).

[85] M. C. Gandikota, K. Pogoda, A. Van Oosten, T. A. Engstrom, A. E. Patteson, P. A. Janmey, and J. M. Schwarz, Soft Matter 16, 4389 (2020), arXiv:1908.03725.

[86] H.-R. Thiam, P. Vargas, N. Carpi, C. L. Crespo, M. Raab, E. Terriac, M. C. King, J. Jacobelli, A. S. Alberts, T. Stradal, A.-M. Lennon-Dumenil, and M. Piel, Nature communications 7, 10997 (2016).

[87] V. Venturini, F. Pezzano, F. C. Castro, H. M. Häkkinen, S. Jiménez-Delgado, M. Colomer-Rosell, M. Marro, Q. Tolosa-Ramon, S. Paz-López, M. A. Valverde, J. Weghuber, P. Loza-Alvarez, M. Krieg, S. Wieser, and V. Ruprecht, Science 370(2020), 10.1126/science.aba2644.

[88] A. J. Lomakin, C. J. Cattin, D. Cuvelier, Z. Alraies, M. Molina, G. P. Nader, N. Srivastava, P. J. Saez, J. M. Garcia-Arcos, I. Y. Zhitnyak, A. Bhargava, M. K. Driscoll, E. S. Welf, R. Fiolka, R. J. Petrie, N. S. de Silva, J. M. González-Granado, N. Manel, A. M. Lennon-Duménil, D. J. Müller, and M. Piel, Science 370 (2020), 10.1126/science.aba2894.

[89] R. J. Hawkins, M. Piel, G. Faure-Andre, A. M. Lennon-Dumenil, J. F. Joanny, J. Prost, and R. Voituriez, Physical Review Letters 102, 1 (2009), arXiv:0902.2078.

[90] L. S. Prahl, M. R. Stanslaski, P. Vargas, M. Piel, and D. J. Odde, Biophysical Journal 118, 1709 (2020).

[91] A. Pathak, Physical Biology 15 (2018), 10.1088/1478-3975/aabdcc.

[92] D. B. Brückner, A. Fink, J. O. Rädler, and C. P. Broedersz, arXiv (2019).

[93] B. A. Camley and W. J. Rappel, Physical Review E - Statistical, Nonlinear, and Soft Matter Physics 89, 1 (2014), arXiv:1405.7088.

[94] R. J. Petrie and K. M. Yamada, Trends in Cell Biology 25, 666 (2015).

[95] M. Raab and D. E. Discher, Cytoskeleton 74, 114 (2017).

